# A DC-sensitive video/electrophysiology monitoring unit for long-term continuous study of seizures and seizure-associated spreading depolarization in a rat model

**DOI:** 10.1101/2025.02.04.635811

**Authors:** Jiayang Liu, Bruce J. Gluckman

## Abstract

There has been a long-term need for a low-cost, highly efficient, and high-fidelity epilepsy monitoring unit (EMU) suitable for home-cage monitoring of small-animal models of epilepsy. We show an accessible, scalable, highly space and energy-efficient EMU capable of fulfilling chronic, continuous synchronized multiple animal monitoring jobs. Each rig within the EMU can provide 16-channel high-fidelity, DC-sensitive biopotential recordings, head acceleration monitoring, synchronized voltammetry applications, and video recording on one freely moving rat. We present the overall EMU architecture design and subsystem details in each recording rig. We demonstrate long-term continuous in vivo recordings of spontaneous seizure and seizure-associated spreading depolarization (SD) from freely moving rats prepared under the tetanus toxin model of temporal lobe epilepsy.

**Significance Statement:** Long-term continuous DC-sensitive biopotential and video recordings are essential for capturing the dynamics of epileptic seizures and seizure-related spreading depolarizations (SD), providing a deeper understanding of their underlying mechanisms. These recordings are invaluable for developing animal models of epilepsy, studying seizure prediction, drug testing, and investigating related neurological conditions such as mental health, aging, and dementia. They also reveal rare phenomena that short-duration recordings might miss. However, traditional methods are resource intensive. The new epilepsy monitoring unit (EMU) introduced in this paper offers a cost-effective and space-saving solution for high-fidelity chronic monitoring of freely moving animals, utilizing compact single-board computers and standard cages without interrupting the recordings.

## Introduction

Epilepsy is the fourth most common neurological disorder in the United States after migraine, stroke, and Alzheimer’s disease (2012). It affects 2.2 million people in the United States and more than 65 million people worldwide (Hirtz, Thurman et al. 2007). Epilepsy is characterized by spontaneous recurrent seizures. Spreading depolarization/depression (SD) is a neurophysiological phenomenon characterized by abrupt changes in intracellular ion gradients, and sustained neuronal depolarization, which lead to loss of electrical activity, altered synaptic architecture, and vascular response. Electrophysiological features of SD recorded in humans closely resemble those observed in animal models (Cozzolino, Marchese et al. 2018). Monitoring SD for research and neuro-intensive care in stroke and traumatic brain injury patients (Lauritzen, Dreier et al. 2011, Dreier and Reiffurth 2015) poses methodological challenges in clinical neurophysiology (Hartings, Li et al. 2017). SD can induce changes across slow and higher frequency bands and a hallmark of SD is a negative near-DC shift in the milli-Hertz range (<0.05 Hz), reflecting mass breakdown of electrochemical membrane gradients, with amplitudes reaching of 20mV or more (Leao 1947, Collewijn and Harreveld 1966, Hansen and Zeuthen 1981).

Long-term continuous biopotential and video recordings are crucial to capture and enhance the study of epileptic seizures. Chronic continuous recordings of small freely-moving animals are important to adequately answer epilepsy-related questions such as the development of animal models of epilepsy (Bertram and Cornett 1993, JEFFERYS 2006, Raedt, Van Dycke et al. 2009), understanding epilepsy physiological mechanism (Walter, Hodge and Hutchinson 1951, Worrell, Gardner et al. 2008, Jiruska, Finnerty et al. 2010), the study of seizure prediction and forecasting (Cook, O’Brien et al. 2013, Kingwell 2013, Wang, Han et al. 2021, Xiong, Nurse et al. 2021, Ibrahim, Emara et al. 2022, Liang, Liu et al. 2022), drug testing, and elucidating mechanisms of epileptogenesis and epileptic maturation (Watanabe, Saito et al. 2017). Moreover, long-term continuous recordings with high resolution and wide dynamic range can be applied for investigating SD, pharmacology and epidemiology, trauma brain injury (TBI), Sudden Unexpected Death in Epilepsy (SUDEP), and comorbidity and interaction with other neurological diseases, e.g., mental health, aging, dementia/AD/PD, and sleep disorders. Importantly, long-term continuous recordings help capture the dynamics of epilepsy that have associated rare events, revealing phenomena that cannot be appreciated with short-duration recordings. The need for exhaustive continuous recording can also be understood by considering that the likelihood of a false negative diagnosis of epilepsy increases inversely with the base seizure rate, yet the ability to do such classification, and to identify the mechanisms surrounding changes in seizure susceptibility, are critical for the development of clinically relevant treatments. Long-term, continuous biopotential and video recordings on multiple animals are typically resource intensive: take sizable space and multiple computers. In this paper, we introduce a new epilepsy monitoring unit (EMU) which is capable of providing a high-fidelity chronic continuous home-cage monitoring of freely-moving animals in a low-cost and space-saving way. We use standard autoclave-ready cages including animal handling, food/water, and cage cleaning without interrupting recording. We use compact single-board computers attached to each cage to acquire video and electrophysiology data and stream data directly to a network-attached storage.

## Materials and Methods

The EMU schematic is shown in Figure 1. Within each rig, through a USB cable and a commutator and isolation system, a DC-sensitive recording system is connected to a single-board computer which is fixed on the cage dome. The single-board computer continuously acquires data (e.g., local field potential (LFP), electrocardiogram (ECG), head acceleration) to network-attached storage (NAS) through an Ethernet cable and a power over Ethernet (POE) supported switch. A separate low-light level-compatible camera system which is also fixed on the cage dome spools video recordings to the same NAS through another Ethernet cable and the same switch. All single-board computers are time-synchronized and communicate with the control station sitting outside the animal facility. Each rig fits into a standard commercial ventilated cage rack with a standard autoclave-ready cage base, which means the EMU can be directly adapted without extra components. To assist with husbandry and cage sanitation, while minimizing direct animal contact, we have further implemented a touch-free cage cleaning method that does not interrupt recordings, which is especially important for epileptic animals that tend to be skittish.

**Figure 1.**
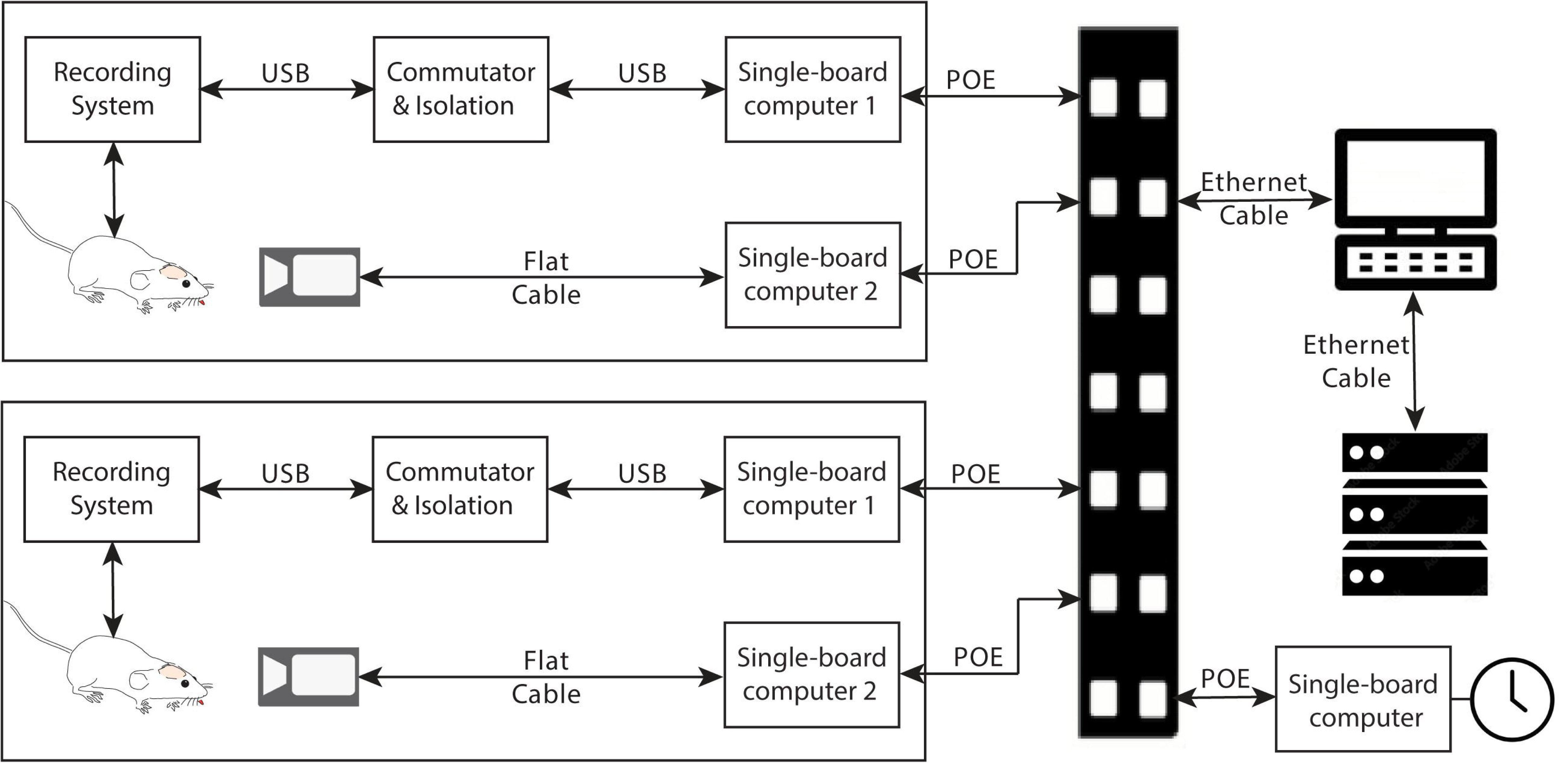
EMU schematic. In each rig, a recording system mounted on the rat’s head is connected to a single-board computer via a USB cable and a commutator & isolation system. A low-light level-compatible camera is connected to another single-board computer. Both single-board computers are fixed to the cage dome and continuously acquire data to a network-attached storage (NAS) through Power over Ethernet (PoE) cables and a switch. A time server is used for synchronization across all rigs.

### Recording System Hardware Design

The current design is adapted from earlier high-performance, low cost, DC-sensitive recording systems suitable for recording from human beings (Jain, Kim and Gluckman 2011) or chronically from mice (Ssentongo, Robuccio et al. 2017) that utilize commercial off-the-shelf (COS) integrated components.

In this paper, we demonstrate a design for chronic freely moving rat recordings. The system has 16 DC-sensitive recording channels, an accelerometer for head acceleration recording; and an electrochemical instrument (EI). Sub-systems arrangement within the recording system has also been improved into a three-layer stack structure and can fit into a 3D-printed box (1 inch × 1 inch × 1 inch). This box is small and light enough to be fixed on a rat’s head as shown in Figure 2. From top to bottom, the three layers are the master board, the daughter board, and the electrode interface board (EIB). The three boards connect through interconnects: the master board and daughter board are soldered together, and this pair can be plugged in or off from the EIB. This design greatly facilitates the experimental implantation and gives flexibility for different recording scenarios.

**Figure 2.**
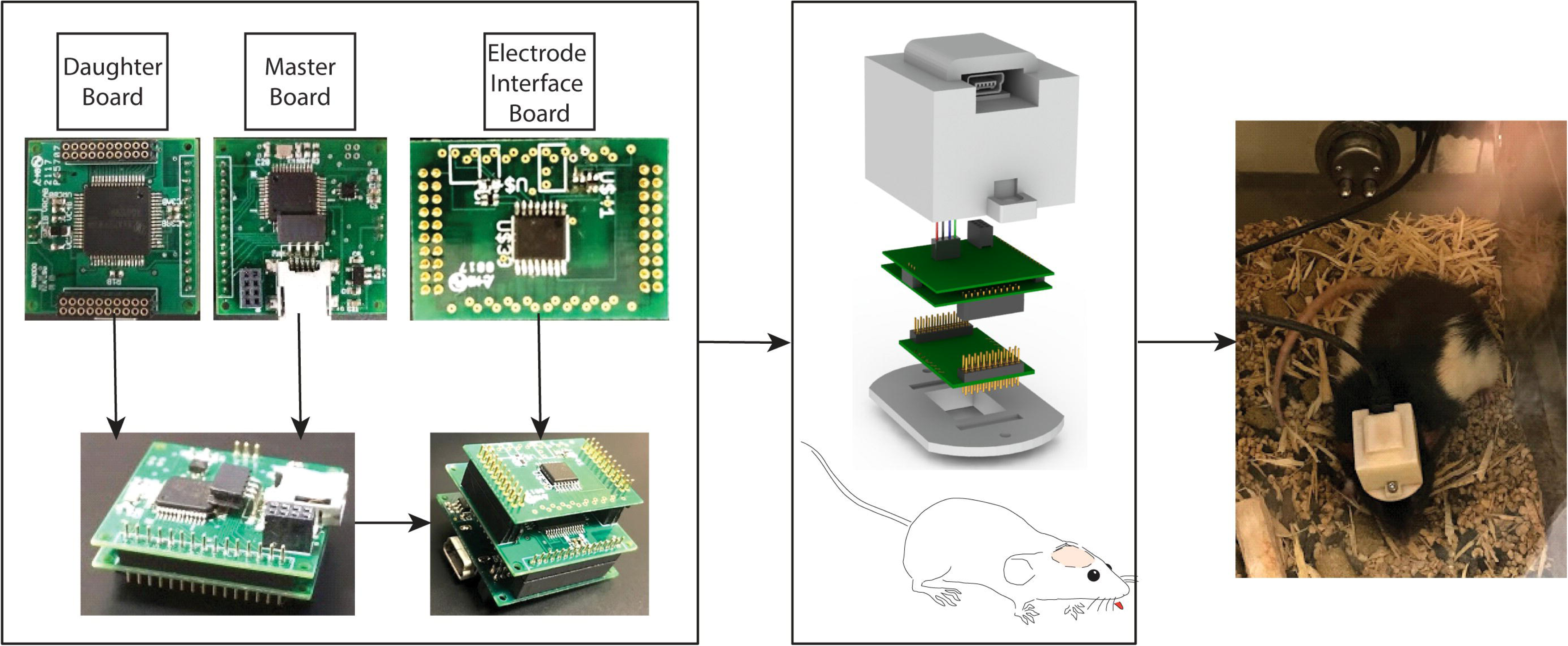
Recording system arrangement. The recording system consists of a master board, a daughter board, and an electrode interface board (EIB), arranged from top to bottom. A 3D-printed box with a cover and a plate holds the three boards. The cover includes a mini-USB connector for connecting the USB cable. The plate secures the EIB and will be cemented to the animal’s head during the surgical process. The master board and daughter board are soldered together and can be plugged in or removed from the EIB via interconnectors.

The master board contains a microcontroller, power regulation subsystem, two system-embedded peripherals (a tri-axis accelerometer and a digital-to-analog converter (DAC)), and a bus to connect to additional peripherals (analog front-end (AFE) amplifiers) on the daughter board.

The micro-controller (Texas Instruments MSP430 series) is responsible for writing to or reading from these peripherals, and communicating with the controlling computer through a USB connection. Biopotential recordings are accomplished through two AFE amplifiers (Texas Instruments, *ADS1299*). The ADS1299 is a high-performance 8-channel, simultaneous-sampling, 24-bit continuous-time delta-sigma analog-to-digital converter (ADC) with a built-in programmable gain amplifier (PGA) featuring low noise and high input impedance. The recording system thus can provide 16 channels of biopotential recordings with sub-µV digitization resolution and ±4.5V dynamic range. Head acceleration is measured using the accelerometer (STMicroelectronics *LIS2HH12*) as a complementary signal for behavioral state of vigilance (SOV) scoring (Sunderam, Chernyy et al. 2007). A digital-to-analog converter (DAC) (Analog Devices *LTC2642*) is included on the master board and serves with the microcontroller as a function generator for peripheral control. The power regulation subsystem is built on the master board. It receives +5V input through the USB connection and supplies regulated +3.3V for digital and +2.5V/-2.5 V for analog power.

The EIB behaves as an interface between implanted electrodes and the recording system. Electrode leads are connected to the EIB in a riveting way using electrode attachment pins, which provide an easy and very reliable long-term connection (Neuralynx, Inc.). The EIB can be plugged/unplugged from the system through interconnectors. In practice, the EIB is permanently cemented to the animal head during surgery allowing the master & daughter boards to be plugged in and connected to all electrodes. After recording, the EIB can be disposed and the master & daughter boards can be reused.

A floating three-electrode potentiostat circuit is built on the EIB as shown in Figure 3. Classic potentiostat designs lock the working electrode at zero potential (ground) of the power supply and drive current to the counter electrode to force the reference electrode to some programed value. Here, in the floating potentiostat design, the reference electrode potential is allowed to float with respect to the amplifier power supply, and current is driven through the working electrode to enforce its potential with respect to the reference electrode. The combination of the function generator on the master board, the three-electrode potentiostat, and appropriate connections to electrodes produces an electrochemical instrument (EI) suitable for a range of electrochemical applications.

**Figure 3.**
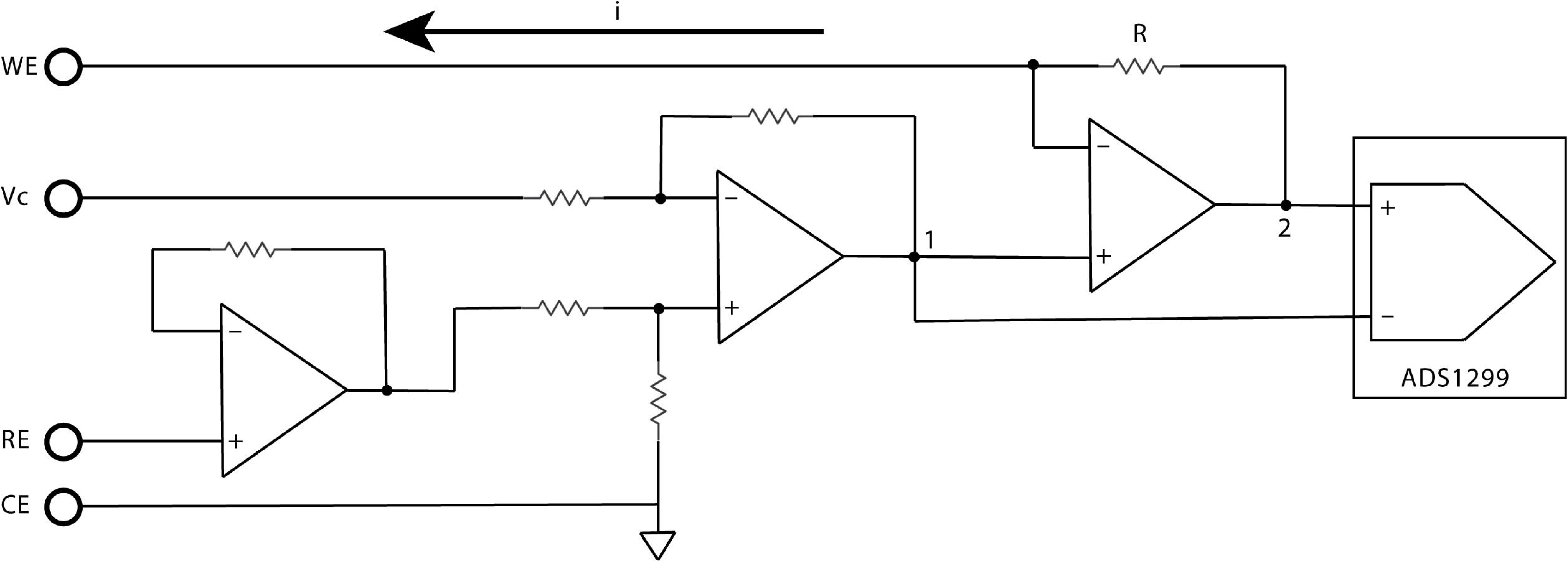
Floating three-electrode potentiostat circuit. Comparing to classic potentiostat setup, the reference electrode potential is allowed to float, and the current is driven through the working electrode to enforce its potential relative to the reference electrode. The current, (*i*), is calculated using the potential difference between points 1 and 2, divided by the resistance (*R*). The potential is recorded by one channel of the ADS1299.

### Recording System Firmware and Software Design

The embedded firmware in the recording system is programmed using program language C and is based on the MSP Driver Library and USB API. The firmware incorporates several functional blocks as shown in Figure 4 and is downloaded into the micro-controller through a custom-made JTAG port on the master board using MSP430 USB-Debug-Interface MSP-FET430UIF Programmer Debugger Emulator.

**Figure 4.**
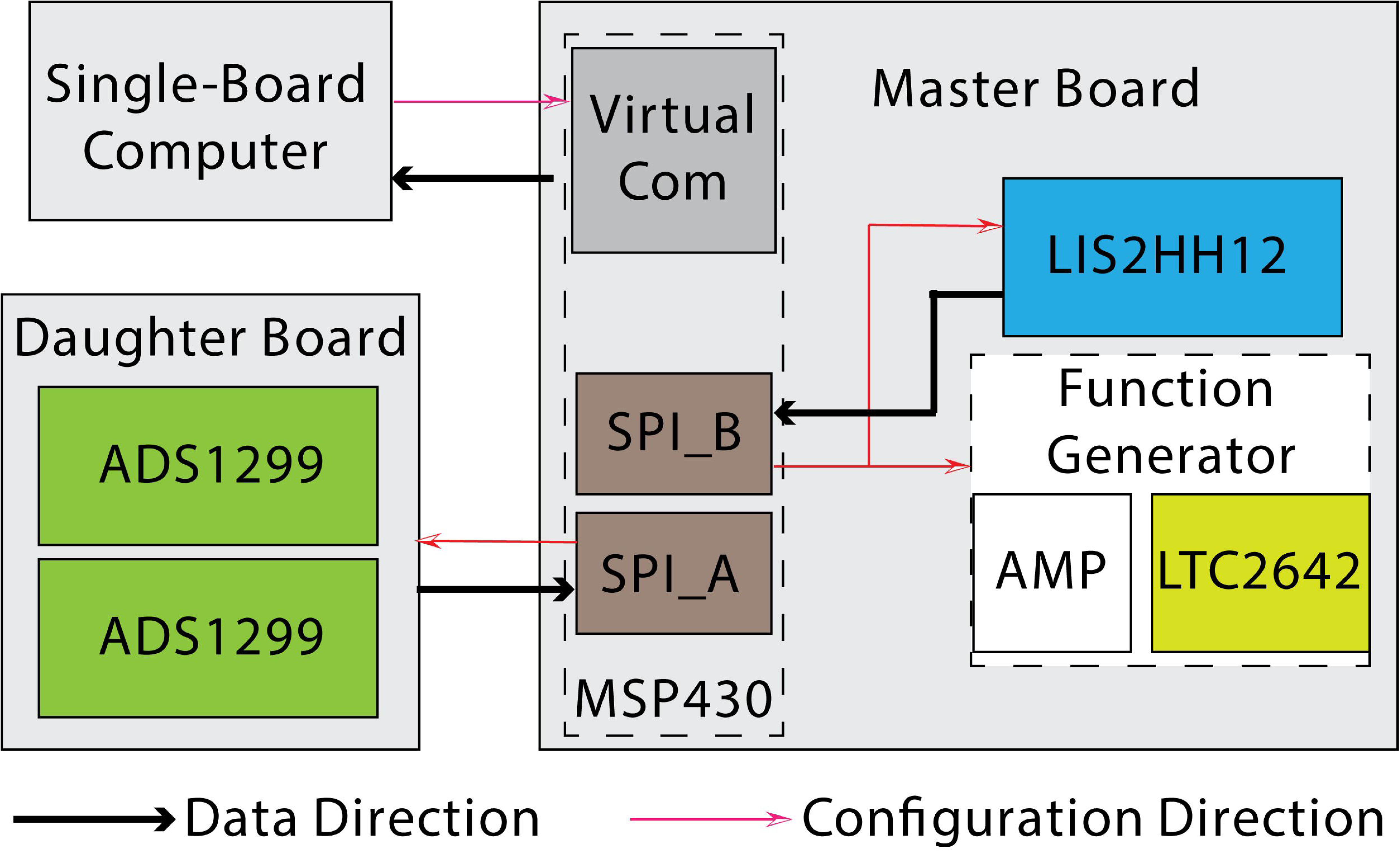
Recording system firmware design. In the firmware design, the microcontroller (MSP430) interfaces with peripheral elements through the universal serial communication interface set to synchronous peripheral interface (SPI) mode. Specifically, it uses SPI_A with the AFEs (ADS1299) and SPI_B with the accelerometer (LIS2HH12) and the DAC (LTC2642). The microcontroller communicates with the single-board computer via the Texas Instruments USB to virtual COM port. The microcontroller receives commands from and sends recorded data packets to the single-board computer through a USB connection.

The micro-controller (*MSP430*) interfaces with the AFEs (*ADS1299*), the accelerometer (*LIS2HH12*), and the DAC (*LTC2642*) through the universal serial communication interface setting in synchronous peripheral interface (SPI) mode. The SPI is a full-duplex interface. Data from two *ADS1299s* and *LIS2HH12* is simultaneously transmitted to the *MSP430*, while configuration inputs from *MSP430* are received by *ADS1299*, *LIS2HH12*, and *LTC2642*, allowing for uninterrupted recording updates. The microcontroller communicates with the single-board computer via USB using the Texas Instruments USB to virtual COM port driver firmware. The microcontroller continuously receives commands from and sends recorded data packets to the single-board computer through the USB connection. Each data packet consists of 63 bytes including 3 bytes-counter, 3 bytes-status of *ADS1299#1*, 24 bytes-*ADS1299#1*’s 8 channels data, 3 bytes-status of *ADS1299#2*, 24 bytes-*ADS1299#2*’s 8 channels data, 6 bytes *LIS2HH12’s* acceleration data. Data stream sending operation occurs in the “background” whenever the USB bus is available until it’s completed to take full advantage of background operation. The function generator in EI is implemented based on a DAC, *LTC2642*. We have implemented five waveforms for different voltammetry applications: constant potential amperometry, fast-scan cyclic voltammetry, long-term pulse voltammetry, differential pulse voltammetry, and sinusoidal coupled with DC potential.

The data acquisition and process software in the single-board computer is coded using Python under the Linux system including recording system configuration, data reading and processing, and data sending to the network-attached storage (NAS) as shown in Figure 5. Input configuration details can be edited at the control station. Through the Ethernet cable, the data is sent to the NAS where it is stored in a binary file with each file started every hour.

**Figure 5.**
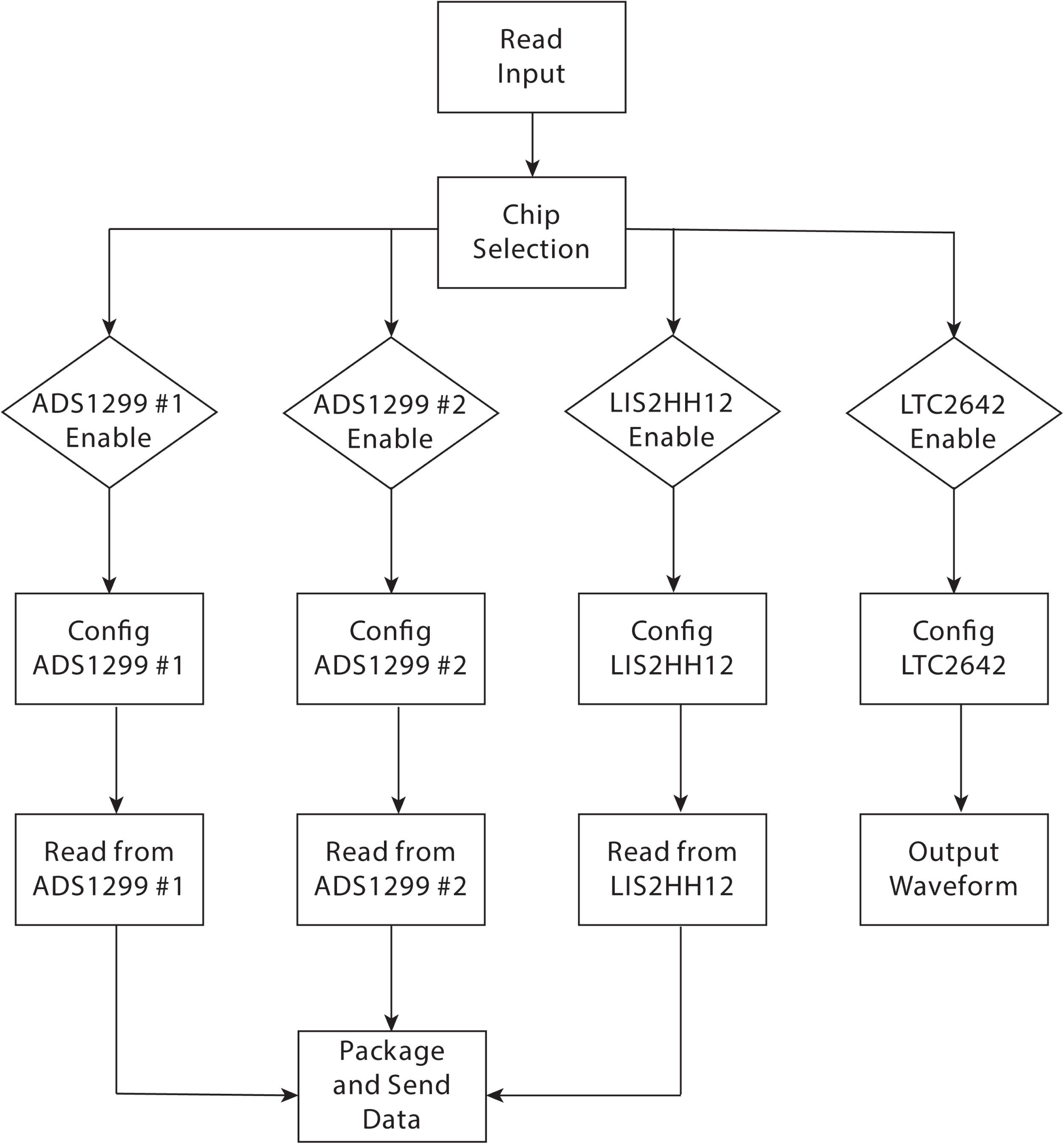
Data acquisition and process software. The data acquisition and processing software on the single-board computer is coded in Python under Linux. It includes recording system configuration, data reading and processing, and data transmission to the network-attached storage (NAS). The input file contains configuration details for all peripheral elements, including two ADS1299 AFEs, the accelerometer (LIS2HH12), and the DAC (LTC2642). These configuration details can be edited at the control station. The recorded data packets are continuously sent to the single-board computer. Each data packet consists of 63 bytes, including a 3-byte counter, 3 bytes for the status of ADS1299#1, 24 bytes for ADS1299#1’s 8 channels of data, 3 bytes for the status of ADS1299#2, 24 bytes for ADS1299#2’s 8 channels of data, and 6 bytes for the LIS2HH12’s acceleration data.

### Commutator and isolation system

Each rig within the EMU is capable of doing chronic continuous recording from a freely moving rat using a commutator and isolation system connected to the recording system through a USB cable. As shown in Figure 6, the commutator provides a USB 2.0 connection and is built with a slip ring (Moog’s Components Group) and a thrust ball bearing (McMaster-CARR 60715K11). The thrust ball bearing, glued with the slip ring box, <u>supports the axial forces</u> from the USB cable weight or various movements from the animal. This design protects the slip ring commutator whose function is sensitive to misalignment and whose bearings are designed to support transverse forces only. The slip ring has six 2A/120VAC circuits and we use 4 of them for USB connection. A 3D-printed mechanical supporter is used to hold both the slip ring and the thrust ball bearing. The whole structure fits in a box. The USB supporter eliminates the vertical tension to wires.

**Figure 6.**
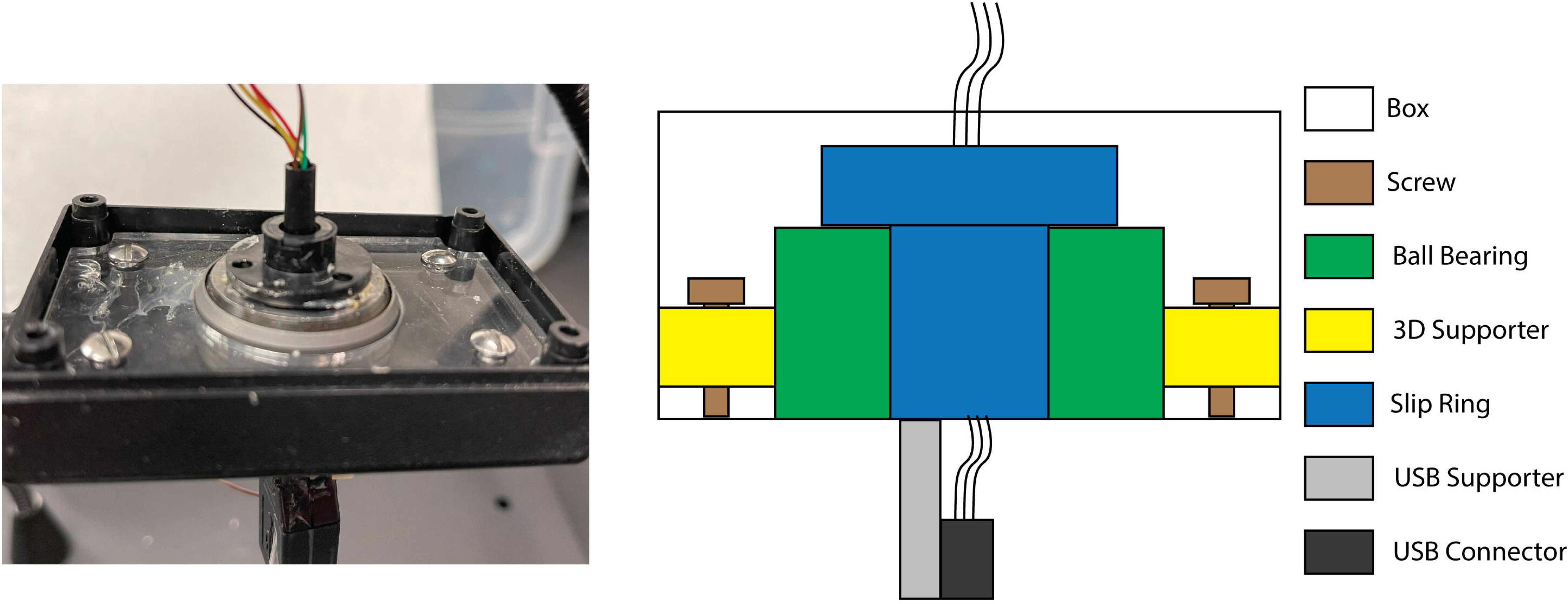
Commutator design. The commutator provides a USB 2.0 connection to the recording system and primarily consists of a slip ring (Moog’s Components Group) and a thrust ball bearing (McMaster-CARR 60715K11). The thrust ball bearing is glued to the slip ring. A 3D-printed mechanical support holds both the slip ring and the thrust ball bearing, and is fixed in a box.

The recording system not only communicates with the single-board computer but also is powered by the single-board computer. The recording system sits on the animal’s head and thus also electrically contacts the animal, which potentially exposes the animal to the risk of electrical shock, burns, and damage directly due to leakage current resulting from improper grounding and electrical isolation (Yeo Siok Been 2007). To alleviate these, we utilize an electrical isolation system which importantly minimizes transmission of 120V/60Hz to the animal. The isolation system helps to minimize pickup of line noise by minimizing the charge needed to tightly couple the amplifier power supply and the subject, which is mediated by current passing through the ground electrode, and therefore minimizes common mode signals. The isolation system incorporates a full/low-speed 5kV USB2.0 compatible digital isolation chip and a 5kV isolated DC-to-DC power converter (Analog Devices *ADuM4160* and *ADuM6000*). The *ADuM4160* provides mechanisms for detecting the direction of the USB data flow and controlling the state of the output buffers. The design of this system meets both creepage and clearance distance criteria in the reinforced isolation type defined in *IEC 60601-1* Standard IEC (International Electrotechnical Commission) 60601-1 (Yeo Siok Been 2007) which defines medical-equipment electrical-safety conditions necessary to protect patients, operators, and the surroundings.

### Video recording system

In addition to biopotential and acceleration recordings, 24-hour continuous video recording of epileptic animals offers behavioral assessment during seizure events. We built a low-light-level compatible video recording system as shown in Figure 7. The video recording system includes a 5-megapixel, fixed focus lens infrared night vision surveillance camera which can provide 1080p at 30 fps video recordings, a 3 Watts infrared light lamp that enables a 24-hour continuous recording, a mechanical supporting structure that can be adjusted to change the video recording coverage, and a single-board computer for video parameters configuration and video file recordings. The camera is connected to the single-board computer through a flexible flat cable. The single-board computer, fixed on the cage dome, continuously sends recorded h264 video files to the NAS, initiating new files hourly.

**Figure 7.**
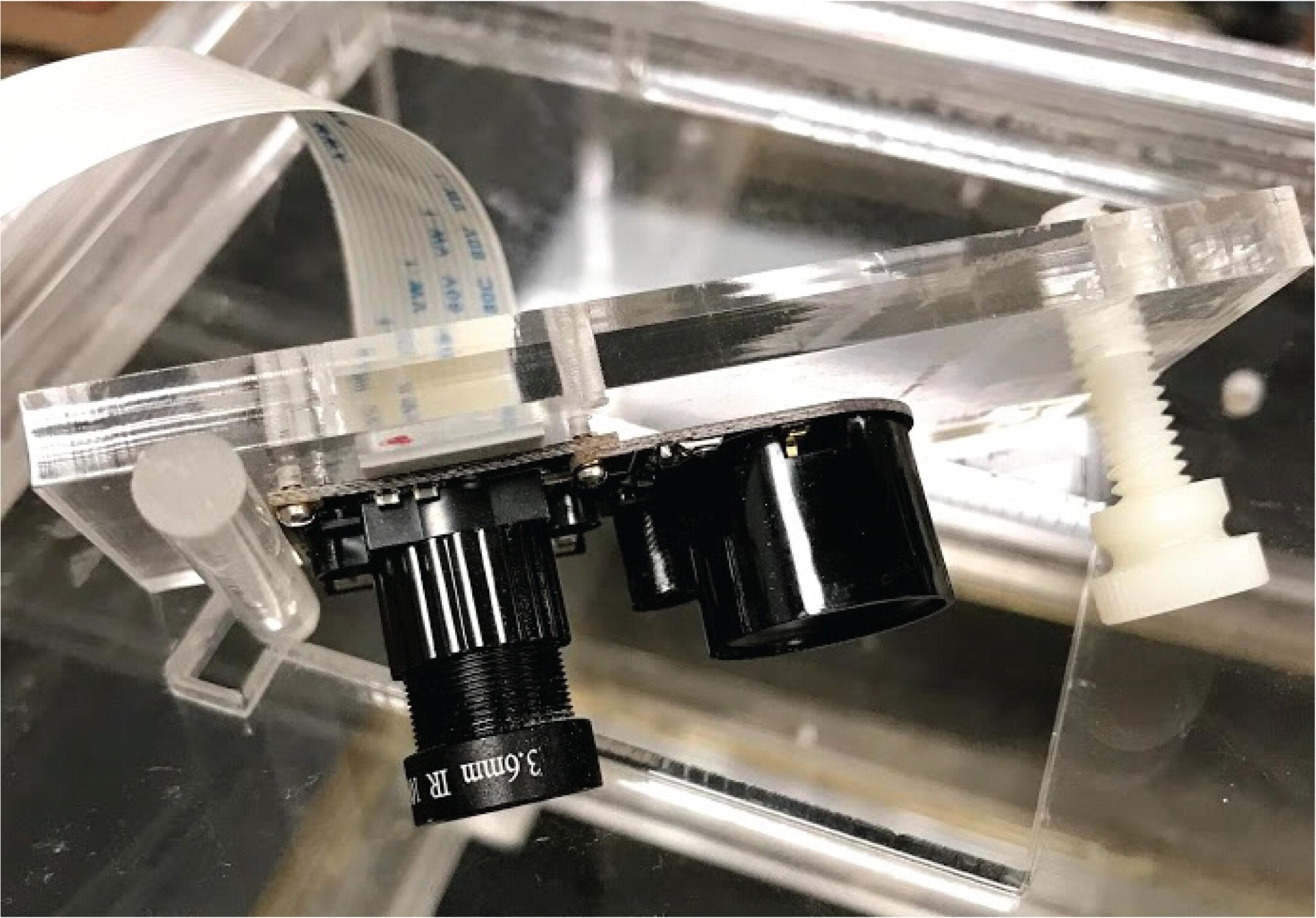
Low-light-level compatible video recording system. The video recording system includes a 5-megapixel, fixed-focus lens infrared night vision surveillance camera capable of providing 1080p video recordings at 30 fps. It also features a 3-watt infrared light lamp that enables 24-hour continuous recording, a mechanical support structure that can be adjusted to change the video recording coverage, and a single-board computer (not shown) for configuring video parameters and recording video files. The camera is connected to the single-board computer through a flexible flat cable.

### Single-board computers and Local Area Network

Commercial products for animal biopotential and video recordings usually involve the use of a desktop or laptop with specifically designed software for data retrieval and processing. In the EMU, we implement one of the most popular single-board computers, Raspberry Pi3 (RPi3). Compared to desktops or laptops, the RPi is low-cost and generates much less heat, and noise. Importantly, it has an extremely small size (3.7”x2.5”x1.0”) and can be easily fixed or removed from the dome of the cage without occupying extra space in the animal room. The RPi3 has four USB ports, an Ethernet port, a Quad Core 1.2GHz Broadcom BCM2837 64bit CPU, and 1GB RAM. It also has a CSI camera port for connecting a Raspberry Pi-compatible camera. <u>All the software and hardware connections are easily adapted to future versions, e.g. RPi5</u>. Each rig can be duplicated for multiple animal monitoring applications. We set up a private local area network (LAN) with a control station (i.e. desktop), a network-attached storage (NAS), (Drobo B810n, Drobo storage company), a POE-supported switch, (NETGEAR-FS726TP, NETGEAR), a time-server and all the other RPi3 used in EMU as shown in Figure 1. On the control station, we can access the data on NAS through the application graphic user interface (GUI) and remotely communicate with each PRi3 through remote applications, e.g., Putty. For a data protection purpose, the LAN is in an “off-line” state, and no RPi3 within the LAN can access the Internet. An additional RPi3 is equipped with a with a real-time clock module, (PCF8523, Adafruit Industries) and set as the time server of the LAN. All other RPi3s are synchronized with this server as time clients through Network Time Protocol. All RPi3s are set using a shell script to follow the same start-up process after being connected with the switch through POE cables. The setup process is to auto-mount on the NAS, get time synchronized with the time server, read in the input file from the control station, auto-start recording data or video, and send data to the NAS.

### Recording on freely moving rats

Initial use of this preclinical EMU was done spanning the years 2017-2020 in the context of two studies whose detailed results are reported elsewhere: one to explore if SD events were associated with seizures and their extent; and the second to investigate tissue oxygenation dynamics during seizures and seizure-associated SD.

Chronic 24/7 continuous bipotential (EEG/ECoG, ECG), head acceleration, and synchronized video recordings were conducted on freely moving rats prepared under the tetanus toxin model (TeTX) of temporal lobe epilepsy (TLE). Tetanus toxin is one of the oldest methods of inducing chronic experimental epilepsies. It was first described by Roux and Borrell (1898) (JEFFERYS 2006). Since then, tetanus toxin has been used to induce seizures in a variety of vertebrate species, but have been utilized mostly in mice and rats (Mellanby, George et al. 1977). Different approaches have been used: hippocampal injections that model focal seizures (temporal lobe seizures), and intracortical injections, which model focal neocortical epilepsy, secondary generalized seizures, and epilepsia partialis continua (Coppola and Moshe 2012). In the last few decades, researchers have worked extensively with tetanus toxin as a model for temporal lobe epilepsy, and have characterized both the mechanisms of action of the toxin as well as seizure development and progression (Jefferys, Borck and Mellanby 1995, JEFFERYS 2006, Sunderam, Chernyy et al. 2006, Sunderam, Chernyy et al. 2009, Sedigh-Sarvestani, Thuku et al. 2014). Usually, after the toxin injection, the animal will start having spontaneously occurring generalized convulsive seizures within 10 days.

### Animal surgery and care

All animal procedures were performed in accordance with the Pennsylvania State University animal care committee’s regulations.

We used Long-Evans rats from Charles River Laboratories, LLC, or ENVIGO, weighing 250 to 450 grams. All rats (Male: 21 Female: 6) were prepared under the tetanus toxin model (TeTX) model of temporal lobe epilepsy (TLE) and the tetanus toxin injection and electrode implantation procedure were described previously (Sedigh-Sarvestani, Thuku et al. 2014). In short, to induce spontaneous epilepsy, 10 to 13 nano-grams of tetanus toxin (Santa Cruz Biotechnology, CAS 676570-37-9) dissolved in 1.3 microliters phosphate-buffered saline (PBS) mixed with 2% bovine serum albumin (BSA) were injected into the rat’s left ventral hippocampus (AP −5.15, ML −5.35, DV −6.1 mm) through a 30-gauge flexible cannula over 15 minutes, which was then left in place for another 30 minutes for tissue relaxation. The recording system can provide up to 16 channels of recordings and based on the experiment’s need, we implant different types and numbers of electrodes. Electrode coordinates were Bregma referenced as shown in Figure 8. 50 µm diameter µRC electrode (Shanmugasundaram and Gluckman 2011) were used as depth electrodes in hippocampus, thalamus, and medial septum. In many cases (especially in later measurements) pairs with tips 125 ∼ 250 µm dorsally apart were used to as bipolar pairs.

**Figure 8.**
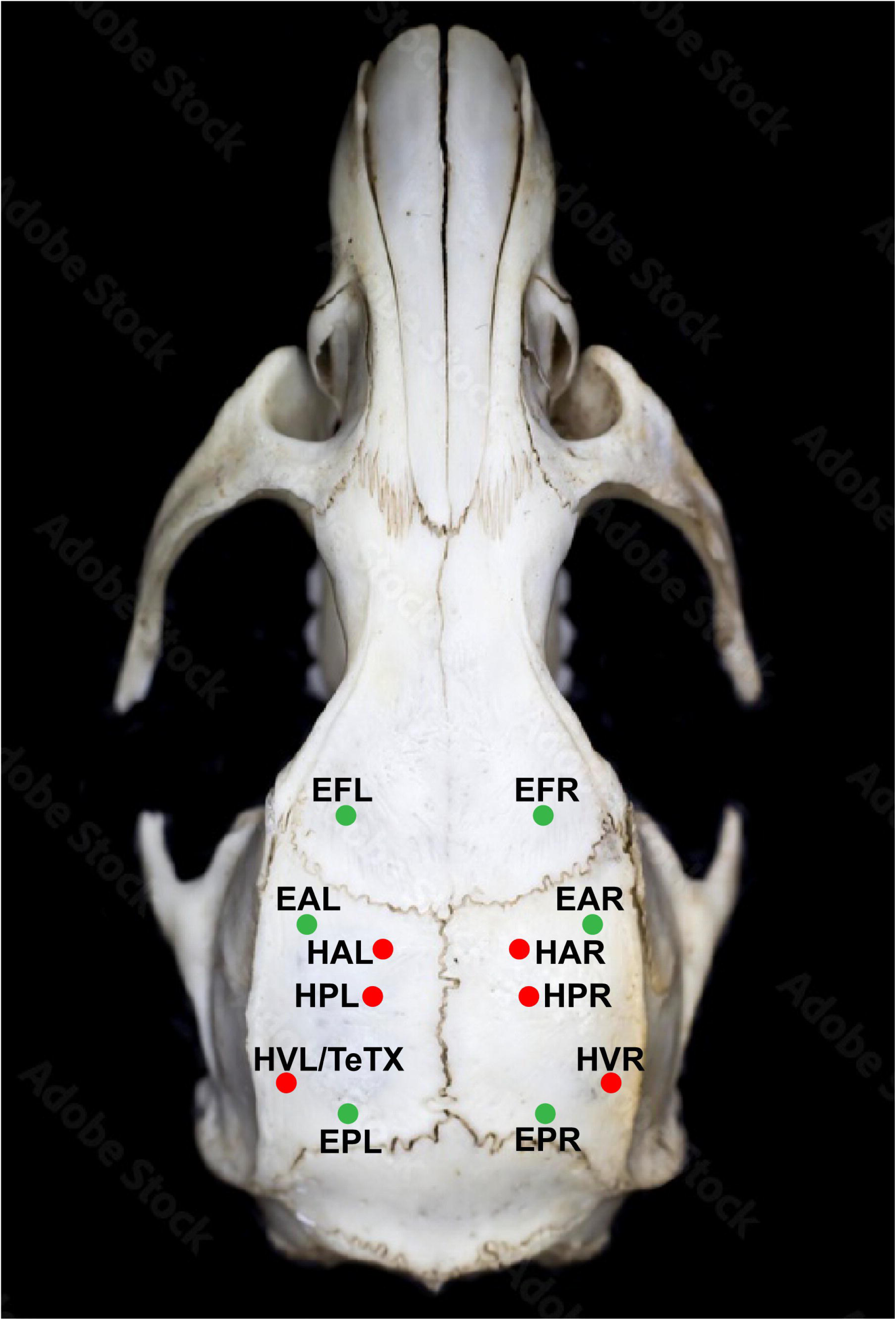
Stereotaxic coordinate of Electrode. All coordinates were Bregma referenced (AP, ML, DV). Red dots show electrodes for differential measurements of the hippocampal local field potentials (LFP). HAL/R (hippocampal anterior left/right): −2.5mm, ±2.0mm, −3.2mm. HPL/R (hippocampal posterior left/right): −3.9mm, ±2.2mm, −2.88mm. HVL/R (hippocampal ventral left/right): −5.51mm, ±5.35mm, −6.1mm. Green dots show stainless steel screws measuring ECoG referentially. EFL/R (EcoG frontal left/right): 2mm, ±3mm. EAL/R (EcoG anterior left/right): 1.5mm, ±4mm. EPL/R (EcoG posterior left/right): −6.5mm, ±4mm. Toxin injection site is at HVL before electrode implantation to minimize additional cortical damage. For oxygen sensing. O2 working electrode is implanted at HAL and reference electrode at HVL after toxin injection.

Specific targets and naming convention included: hippocampus at HAL (hippocampal anterior left), HAR (hippocampal anterior right), HPL (hippocampal posterior left), HPR (hippocampal posterior right), HVL (hippocampal ventral left), HVR (hippocampal ventral right) to provide differential measurements of the hippocampal local field potentials (LFP). Stainless steel screws measuring ECoG referentially are at EFL (EcoG frontal left), EFR (EcoG frontal right), EAL (EcoG anterior left), EAR (EcoG anterior right), EPL (EcoG posterior left), and EPR (EcoG posterior right).

For amperometry based tissue oxygen measurements, a 250 μm diameter Platinum (Pt) wire working electrode was implanted targeting the hippocampus and custom Ag/AgCl reference electrode was implanted into the cortex along the same trajectory as used for toxin injection to minimize additional cortical damage. The Ag/AgCl electrode was made by inserting a 100um diameter Ag wire into a 155um diameter Polyamide tube filled with Ag/AgCl ink (CI-4001 Silver/Silver Chloride/Vinyl, Nagase America LLC). Animal implantation details are listed in Table 1. Note that each animal implantation has slight differences based on the experimental purpose and details have been listed in Supplementary Tables. Animal ECG was monitored simultaneously with three lead electrodes implanted in the precordium. Depth electrodes are secured in place and cortical screw electrodes electrically isolated via dental cement. All leads are connected to the recording system and encapsulated within the 3D-printed head mount. After surgery, rats were returned to individual standard autoclave-ready cages with free access to food and water and maintained at a 12-hour light-dark cycle with lights on between 7 am and 7 pm. We allowed a seven-day post-surgery recovery before cabling initiating recordings. The data sampling rate was 1000Hz. During the continuous data recording periods, rats can move freely in the cage, and food-adding, water-adding, and cage-changing works were done without interrupting the recordings. In Video 1, we demonstrate a technique to change the old cage with a clean one with food and fresh water. In short, after putting two cages side by side and moving the cage dome to the middle, the rat will jump into the new cage by itself while still under recording.

**Table 1.**
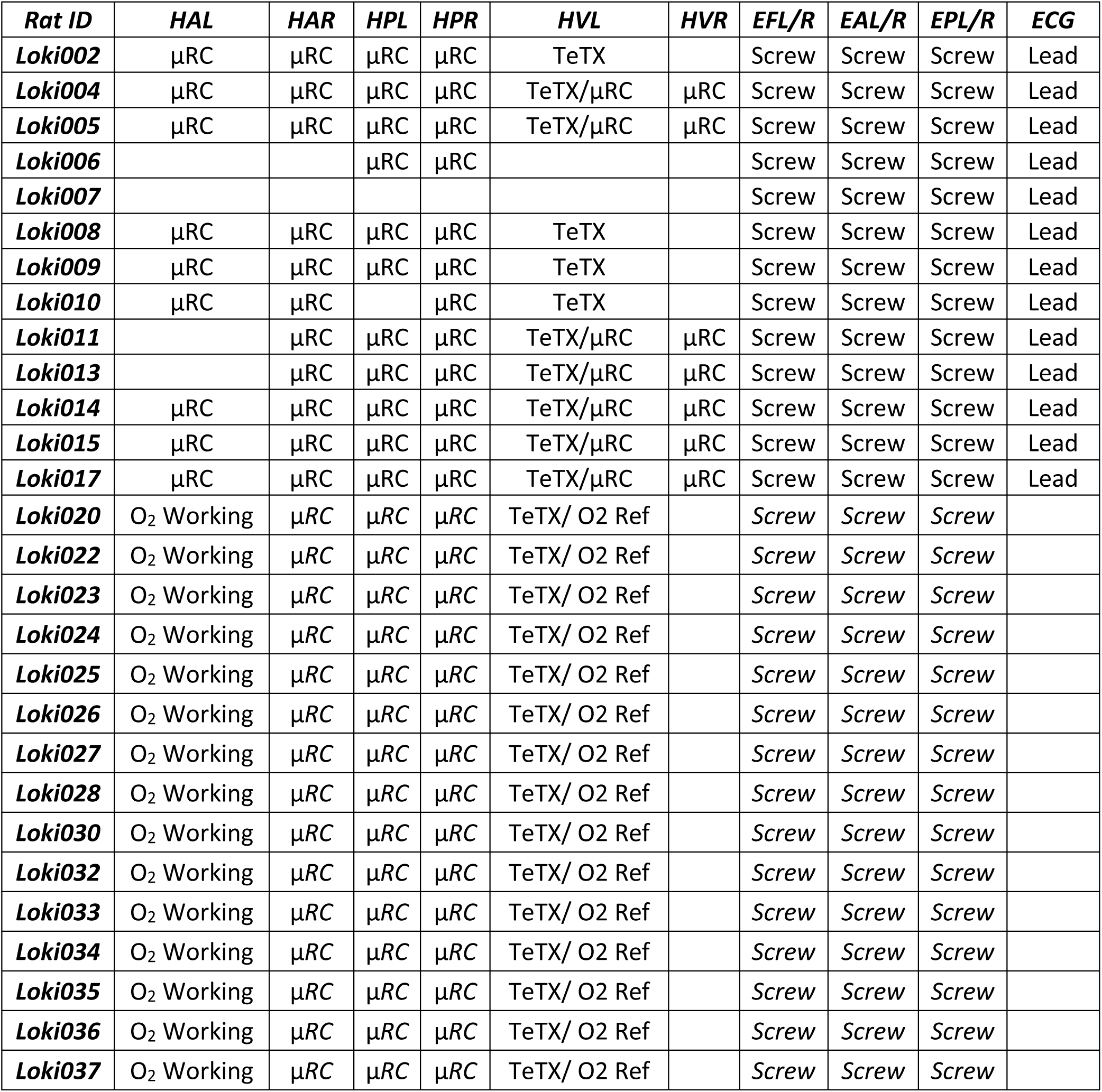
Electrode implantation summary. Hippocampus target: (AP, ML, DV) (mm): HAL (hippocampus anterior left): (−2.5, −2.0, −3.2), HAR (hippocampus anterior right): (−2.5, +2.0, −3.2), HPL (hippocampus posterior left): (−3.9, −2.2, −2.88), HPR (hippocampus posterior right): (−3.9, +2.2, −2.94), HVL (hippocampus ventral left): (−5.51, −5.35, −6.1), HVR (hippocampus ventral right): (−6.0, +5, −5.5). Cortex: (AP, ML) (mm): EFL (ECoG frontal left): (+2, −3), EFR (ECoG frontal right): (+2, +3), EAL (ECoG anterior left): (+1.5, −4), EAR (ECoG anterior right): (+1.5, +4), EPL (ECoG posterior left): (−6.5, −4), EPR (ECoG posterior right): (−6.5, +4). µRC (micro-reaction chamber electrode), O2 Working (Oxygen sensing working electrode, Pt wire), O2 Ref (Oxygen sensing reference electrode), TeTX (Tetanus toxin).

### Data analysis

Recorded data include local field potentials (LFP) from µRC electrodes targeting the hippocampus, ECoG from screw electrodes over the cortex, ECG recordings, three-axis head acceleration signal, and tissue oxygenation signal. The tissue oxygenation signal is the current response from the constant potential amperometry (CPA). Data is processed offline using custom-written MATLAB (MathWorks Inc.) programs for re-referencing, filtering, spectral analysis, and behavior annotation. Unless otherwise indicated, raw ECoG and hippocampal LFP are band-pass filtered at 0.5 ∼ 125Hz to highlight field potentials and seizure dynamics. Averaged power spectrum density (PSD) is calculated with a welch-average, 2048 window after application of the 0.5∼125Hz bandpass filter. Spectrograph and mean square root (MS) power of acceleration are calculated using 1 second window after 2∼125Hz bandpass filtered. **Seizure Detection:** Seizures were detected by a stereotypical increase in spectral power that typically initiates with a sentinel spike followed by a burst of 9-16 Hz hippocampal spikes (Finnerty and Jefferys 2000), spreads through the cortex, and ends with a sharp decrease in spectral power. For practical purposes, we count only events that are longer than 10 s, and spaced apart by at least 10 minutes.

### Spreading Depolarization Detection

SD detections were done from depth electrode measures either differential within bipolar pairs, or referenced to a cortical screw electrode. SD events were detected from signals low-pass filtered with a cutoff at 0.5 Hz, with onset defined by a downward crossing of a 7.5 mV threshold in the signal with respect to the value 3 seconds before. SD offset was then detected by an upward crossing of a threshold defined as 2 mV above the SD onset potential.

### Cardiac RR Interval Measurement

For times when the cardiac signal is free of movement/chest muscle artifact, cardiac ventricular contractions were detected by first band-pass filtering the cardiac signal from 15∼125 Hz, then locally finding the peak times of the R wave. RR intervals were then computed as the first difference in R peak times.

### State of Vigilance Classification

State of vigilance (SOV) was marked as one of the three states: REM state characterized by a spectral peak in the theta (4 ∼ 7) frequency band of hippocampal LFP and by an absence of acceleration except during brief muscle twitches; NREM state characterized by maximal power in the delta (0.5 ∼ 4 Hz) frequency band and by an absence of acceleration; Wake state characterized by the accelerometer activity (Sunderam, Chernyy et al. 2007, Bahari, Kimbugwe et al. 2021). *In vivo* CPA response current was low-pass filtered at 1Hz and then down-sampled from 1kHz to 20Hz.

## Results

Data were collected from 27 TeTX epileptic rats and stored on the NAS (Data summary in Table 2) in the context of two studies whose detailed results are reported elsewhere: one to explore if SD events were associated with seizures and their extent; and the second to investigate tissue oxygenation dynamics during seizures and seizure-associated SD.

**Table 2.**
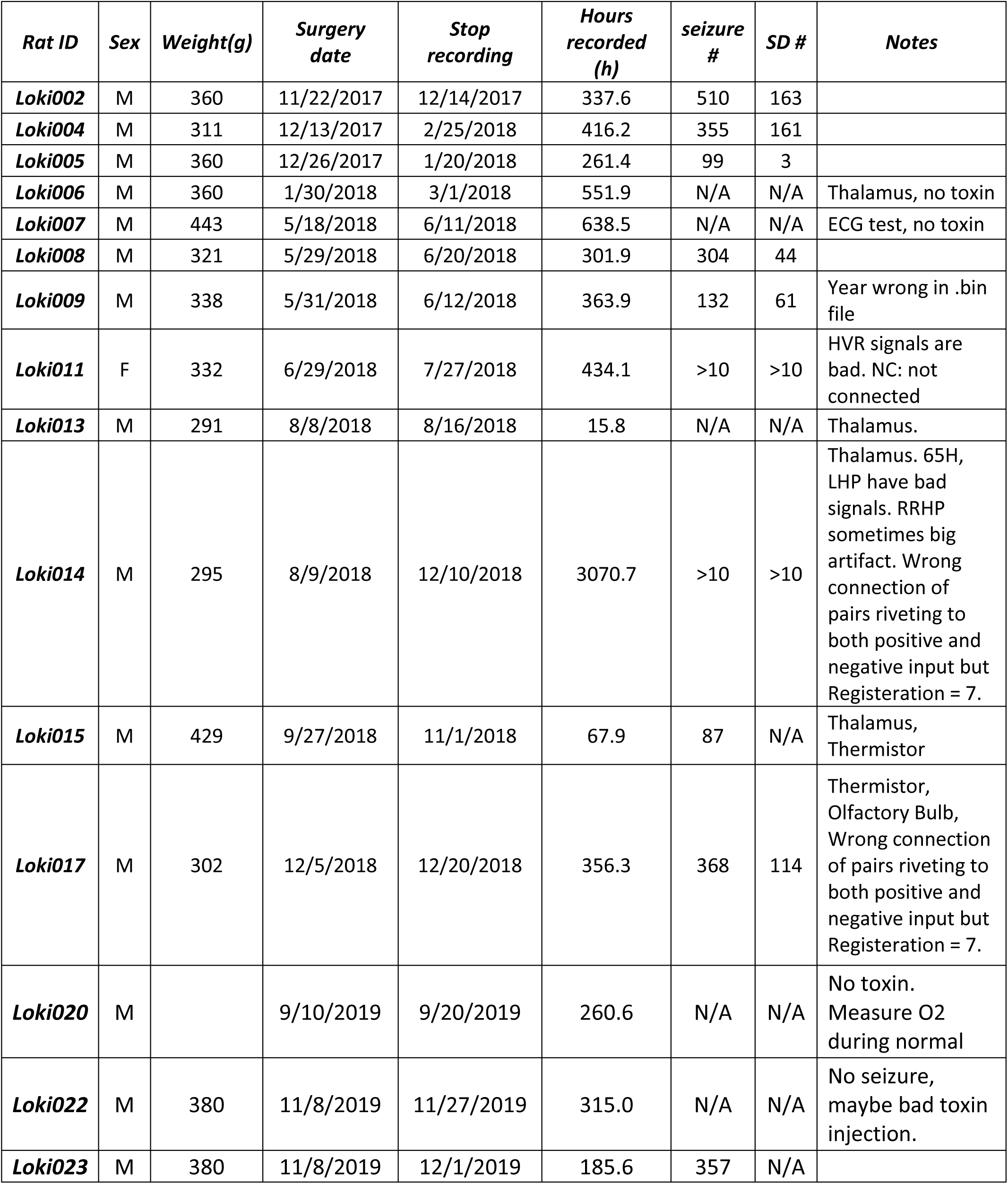
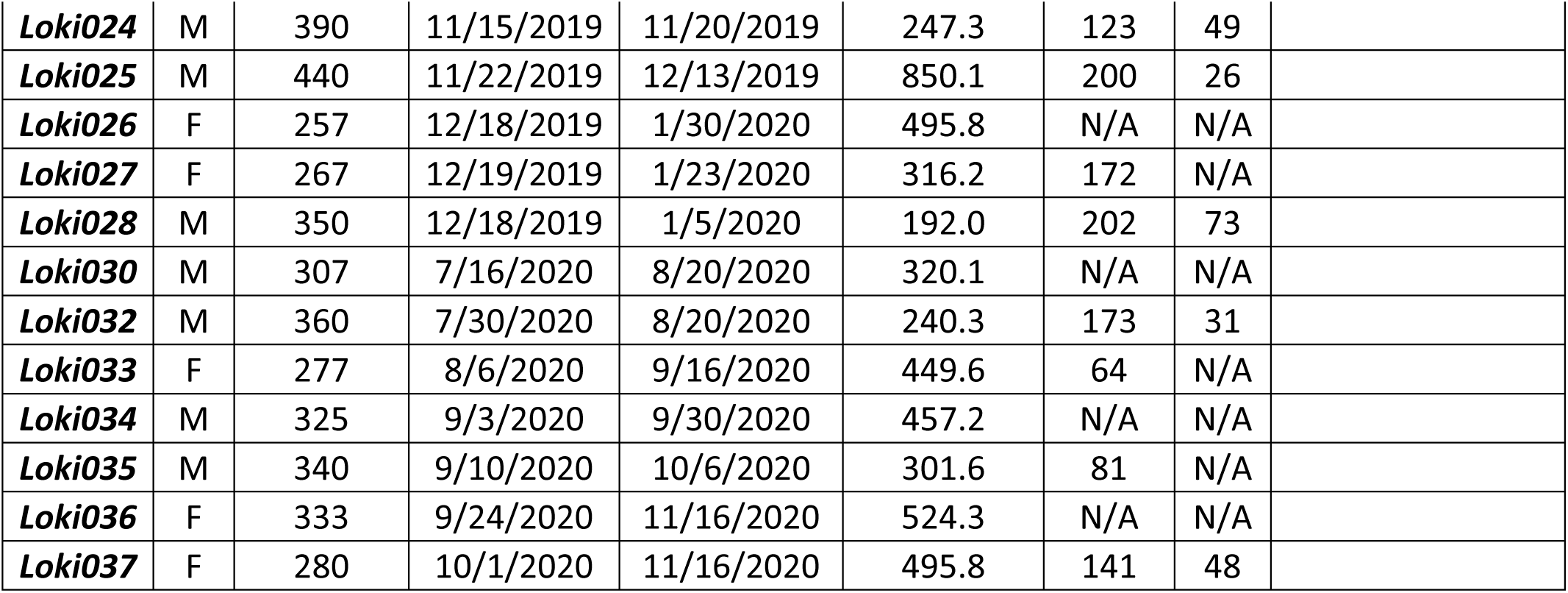
Animal surgical information and data summary.

### Signal Quality from homecage measurements in Freely Moving Rats

The recording system’s high input impedance coupled with the improved impedance µRC electrode provides a high-fidelity signal. In Figure 9a we show the average spectral power four typical electrophysiology channels, computed in 2.048 s windows and averaged over an hour of recording. No 60 Hz line noise is detectable here, and no notch filter was applied anywhere within the signal path. In Figure 9b are shown four 20-second traces of stereotypical normal behavior: exploratory theta, transition from wake to sleep, transition from NREM to REM, and end of REM with brief awakening and movement respectively. Two selected channels spectrograph and the acceleration MS power illustrate the transition clearly (the triangle shows the transition point) in Figure 9c.

**Figure 9.**
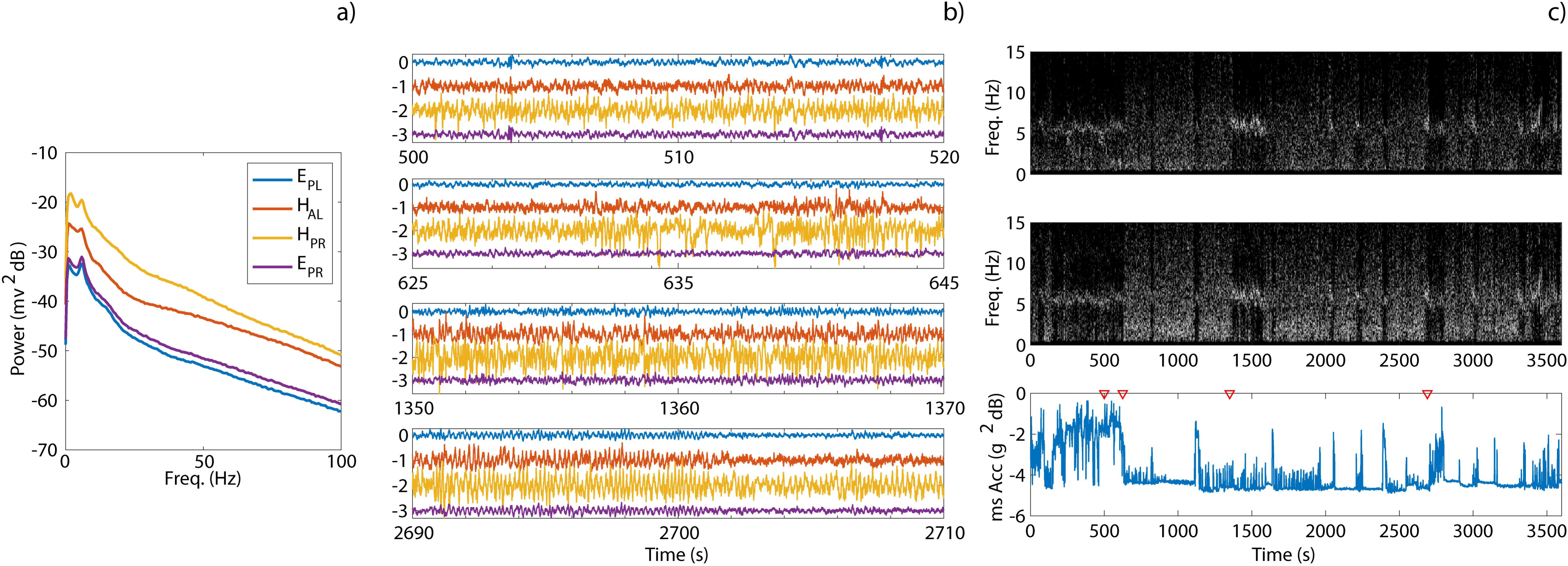
Example recording demonstration. **A)** The averaged power spectral density (PSD) for these traces. **B)** Four 20-second example traces (EPL/R, HAL, HPR) illustrate exploratory theta, the transition from wake to sleep, the transition from NREM to REM, and the end of REM with brief awakening and movement, respectively. There is no 60Hz peak without applying a notch filter. **C)** The spectrograph of two selected channels and the acceleration mean square root power show the transitions (with the triangle indicating the transition point).

### Spontaneous Seizures and Seizure-associated Spreading Depolarization (SD) in Freely Moving Rats

Identifying features of an SD include a large negative shift in tissue potential amplitude of an order of 10 to 30 mV and duration lasting for approximately 30s to 60s (Grafstein 1956, Kraig and Nicholson 1978, Somjen, Aitken et al. 1992). The DC-sensitivity and large dynamic range and bit depth are critical for being able to span the dynamics of normal field potentials, seizures, and spreading depolarization in a single recording. From this data we found frequent instances of spontaneous seizures and seizure-associated SD events. One example of one seizure and seizure-associated spreading depolarization (SD) event is shown in Figure 10. The upper panel of Figure 10 shows band-filtered ECoG, and hippocampal LFP measurements, and the lower panel includes low-pass filtered differential hippocampal LFP measurements. The ictal period is marked yellow. The DC-band recordings of seizure and associated SD are shown in the lower panel of Figure 10. The characteristic DC shift of SD is marked using triangles.

**Figure 10.**
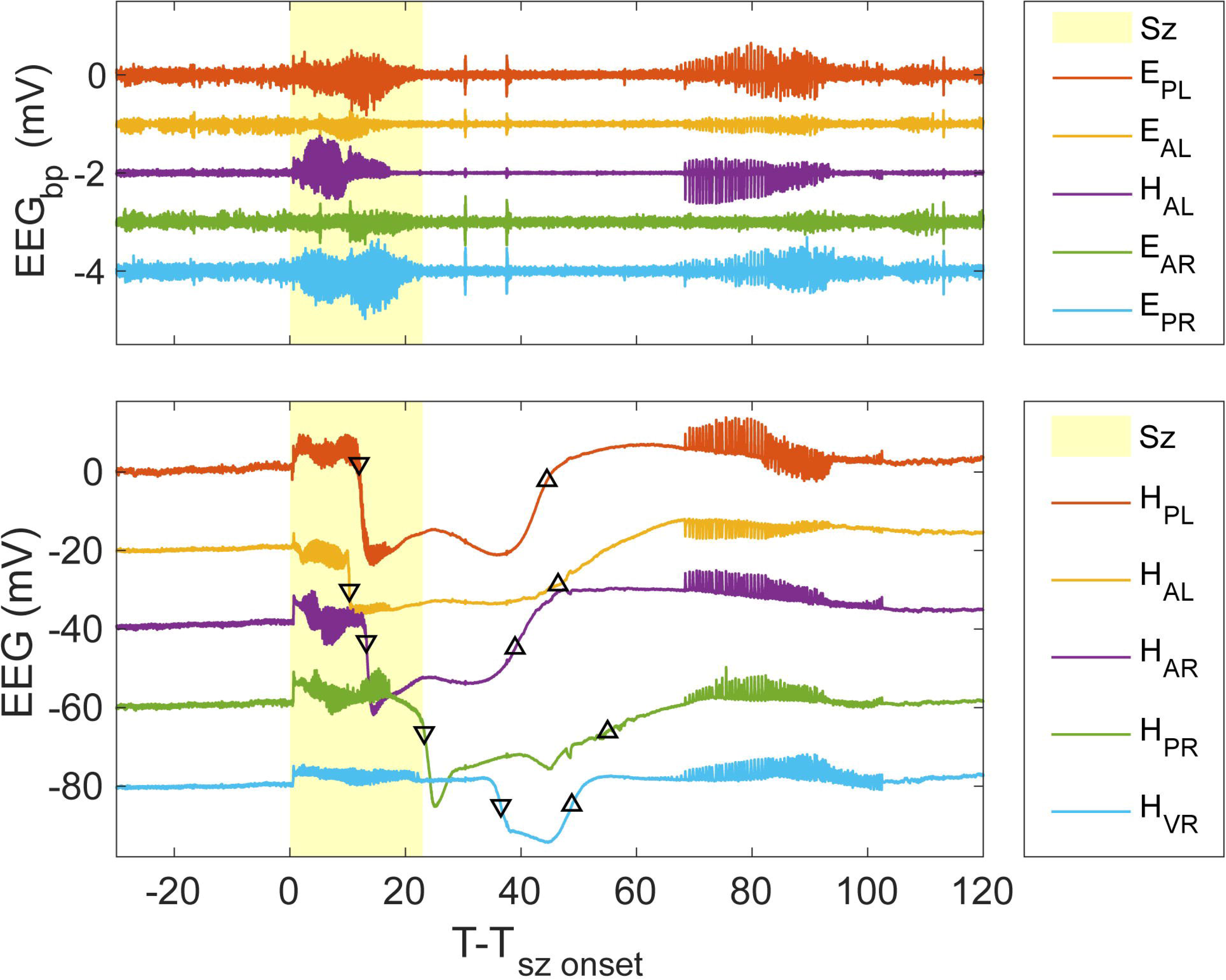
Example of one seizure and seizure-associated spreading depolarization (SD) event. The upper panel shows band-filtered ECoG and hippocampal LFP measurements, and the lower panel displays low-pass filtered differential hippocampal LFP measurements. The ictal period is marked in yellow. The lower pane also shows DC-band recordings of seizures and associated spreading depolarizations (SD), with triangles indicating the SD propagation direction.

These DC measurements clarify that the depressed EEG activity and the later after-discharges are seizure-associated SD events and recovery from it. This phenomenon would not be observable without DC-sensitive recordings. Abnormal fluctuations in brain activity are likely to subtly modify cardiac function long before the microscale instability propagates and results into an overwhelming seizure (Bragin, Wilson and Engel 2000, Schevon, Ng et al. 2008, Stead, Bower et al. 2010). In (Bahari, Ssentongo et al. 2018), the authors demonstrated a brain-heart biomarker for epileptogenesis in a murine mouse model based on occasional/rare long cardiac RR intervals that are preceded by fluctuations in cortical activity weeks to months before the animals’ first seizures. In our TeTX model, from these recordings we find similar signatures in multiple animals ∼4 days before their first convulsive seizure, and only a few such events within a limited period of a few hours. An example of one of these events is example is shown in Figure 11. In the upper panel can be seen an abnormal 7∼8 Hz oscillation which lasts about 5 s and is synchronized across subcortical regions. Approximately 5 s later a cardiac arrhythmia event with substantially long RR intervals occurs (lower panel). In general, the cardiac measures were significantly confounded with muscle and movement artifact so observations of such arrythmia were limited to times when the animal was substantially still, such as when asleep. Complete analysis of such dynamics is left to future studies done with better cardiac electrode design and placement technique.

**Figure 11.**
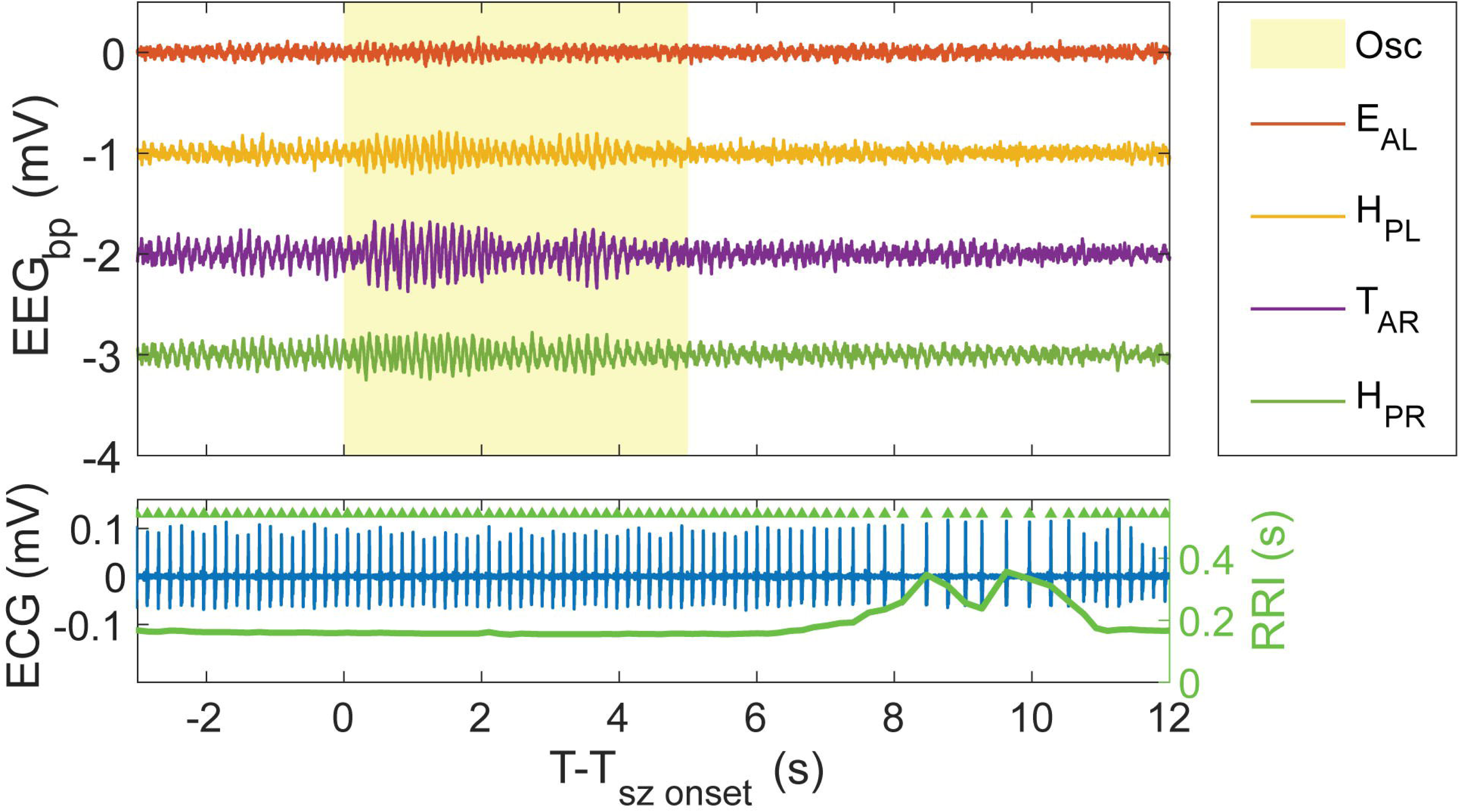
Example of abnormal cortical discharges preceding altered cardiac functions. Generated from Loki011. The upper panel shows band-filtered ECoG and hippocampal/thalamus LFP measurements. TAR: thalamus anterior right. The abnormal 7∼8Hz oscillation is marked in yellow. The lower panel displays low-pass filtered ECG measurements. The heart rate time-series is marked blue and the point-by-point RR interval (RRI) is marked green.

### Oxygen sensing using built-in EI

The built-in EI in each rig extends our EMU’s application scenario, such as constant voltage amperometry (CPA) applications for tissue-level oxygen concentration monitoring. We performed CPA oxygen sensing both *in vitro* for calibration purposes and *in vivo* in freely moving rats to monitor tissue-level oxygenation.

### O2 Calibration

We implement a three-electrode electrochemical cell connecting to our EI. Specifically, the working electrode (WE) is a platinum (Pt) wire, the reference electrode (RE) is an Ag/AgCl pellet, and the counter electrode (CE) is a Pt plate. During the experiment, oxygen content was known by first preparing two separate PBS solutions from the same sample, one was saturated with air, formed by bubbling air through it, and the other with virtually no oxygen formed by bubbling Nitrogen through it. Three electrodes were submerged in the air-saturated solution, and N_2_-saturated solutions were added. The experiment temperature was room temperature and the PBS solution’s pH value was 7.48 (pH Meter, Model P771. Anaheim Scientific). The resulting oxygen concentration is derived from standard stoichiometric calculations. We implemented the CPA measurement by setting the bias potential on the working electrode to −0.65V with respect to the reference Ag/AgCl electrode. Each point came from an addition of a fixed amount of N_2_-saturated PBS solution. Following each addition, we recorded for about 5 minutes, and the reported value was calculated from the middle 90s data of the 5 minutes reject transients associated with the adding N2-saturated PBS solution process. The CPA calibration result is shown in Figure 12. In brain tissue, the oxygen concentration, the partial pressure of oxygen (pO2), and the oxygen tension are mutually related values that, in principle, can be derived from one another. The oxygen concentration is expressed in moles per liter (mol/l) which is usually used for measuring dissolved oxygen. The pO2 reflects the amount of free oxygen molecules and equals to the pressure that oxygen would exert if it occupied the space by itself (Subczynski and Swartz 2005). The calibration is well-approximated by a linear fit over the physiological range from ∼0.09 mM/L (the tissue oxygen concentrations) to ∼0.23 mM/L (the arterial oxygen concentration) (Ortiz-Prado, Dunn et al. 2019) marked by the purple and blue lines.

**Figure 12.**
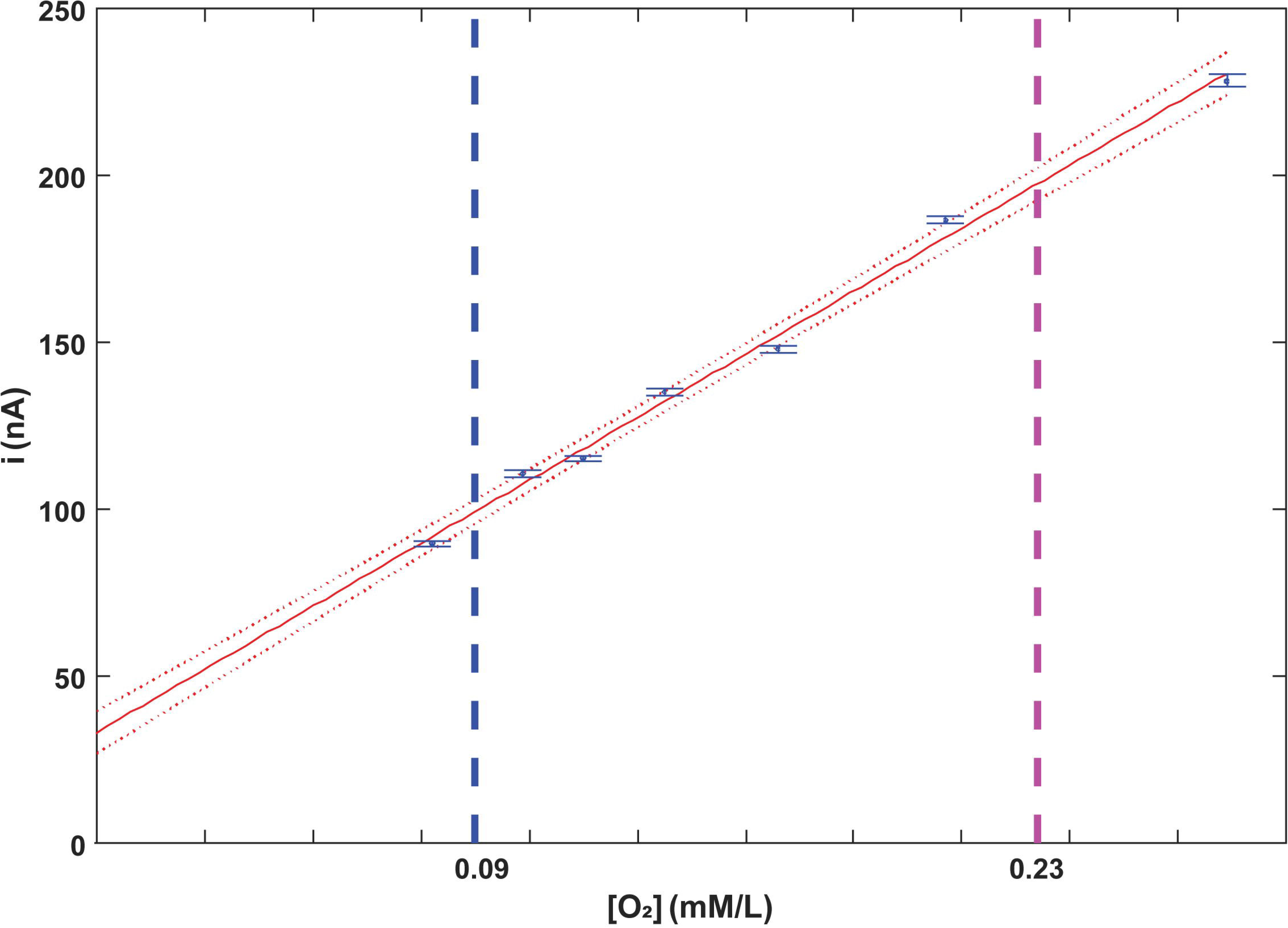
CPA calibration result. A linear fitting result is shown, with each point derived from the addition of N2-saturated PBS solution. For each addition, we recorded for 5 minutes, and each value was calculated from the middle 90 seconds of data to eliminate transients associated with the addition process.

### *In vivo* O2 measurements

We used this system to record brain-tissue oxygen dynamics using CPA from 15 animals (Loki020∼037 in Table1&2) through both normal SOV, seizure, and seizure-associated SD events. Shown in Figure 13 are the tissue-level oxygen dynamics from a single hour of time from one rat (Loki028) using CPA, superimposed on a color-coded hypnogram. This hour was chosen as one without seizures and had multiple sleep/wake cycles. Also shown are the average O2 concentrations within state (red REM n=73, green NREM n=167, blue Wake n=220), as measured across 2 full days, and averaged over SOV bouts that started/ended at least 1 minute from any seizure or SD event. Figure 13 shows a consistent increased CPA measure from NREM to REM transition and a drop of CPA from REM to NREM or Wake transition. This increased CPA during REM Sleep marks a higher oxygen level and may come from an increased cerebral blood flow since REM sleep is characterized by increased neuronal activity, metabolic demand, and cerebral blood flow, leading to greater oxygen delivery. On the other side, a decreased CPA during NREM sleep marks a lower oxygen level since during NREM sleep the animal has reduced neuronal activity and metabolic demand, and cerebral blood flow. Our result of increased constant potential amperometry (CPA) signals during REM sleep and decreased signals during NREM sleep aligns with findings from studies (McHugh, Fillenz et al. 2011, Dash, Tononi and Cirelli 2012) measuring brain tissue oxygen levels in rats using CPA.

**Figure 13.**
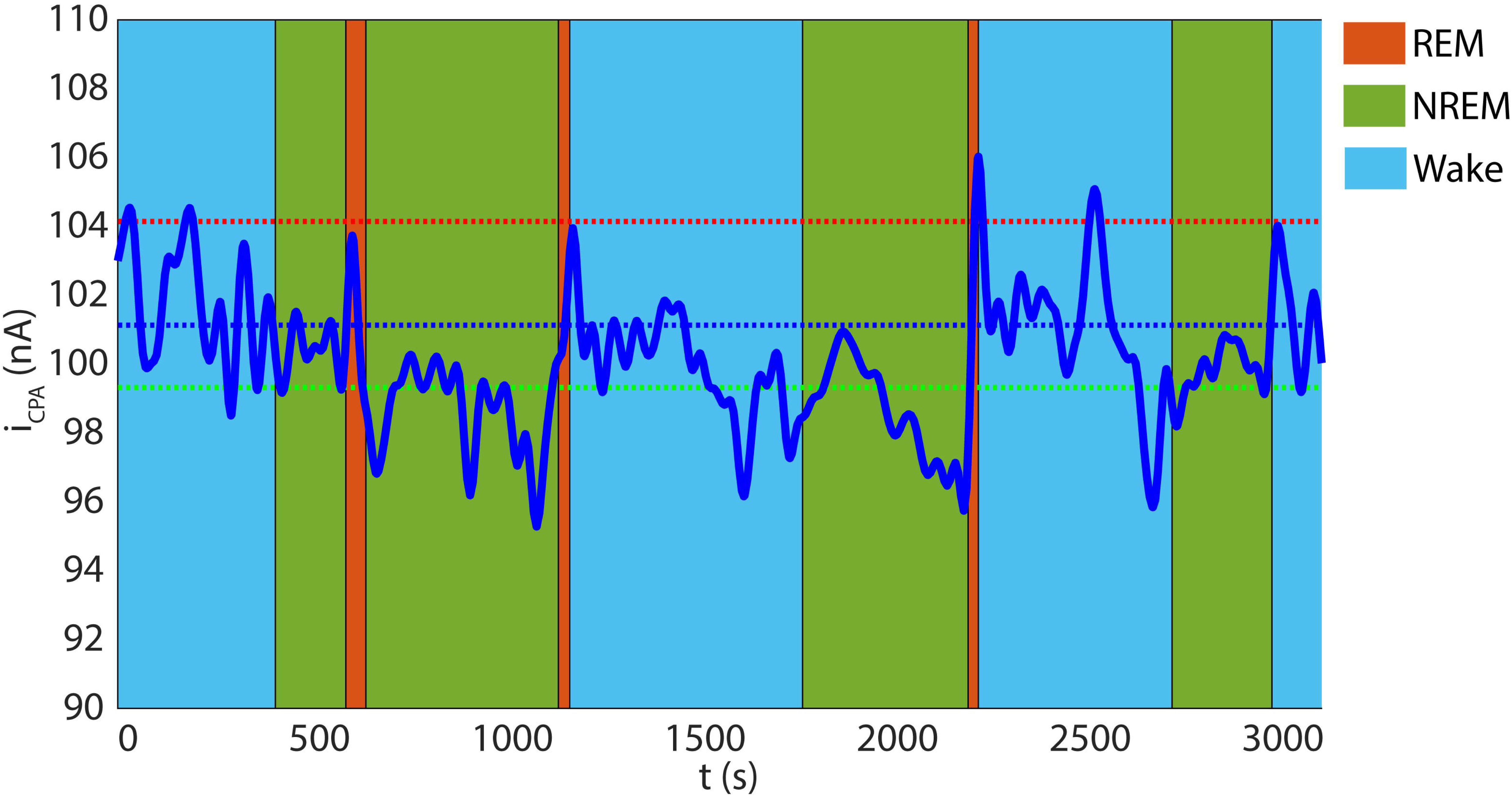
Example of in vivo CPA current recording. One hour out of a two-day CPA current response from one rat (loki028) during a normal SOV. During this period, the animal experienced REM (red), Wake (blue), and NREM (green) states multiple times. The red dotted line represents the mean value of 73 REM state responses over two days of recordings, the green dotted line represents the mean value of 167 NREM state responses, and the blue dotted line represents the mean value of 220 Wake state responses. SOV bouts that started or ended within 1 minute of a seizure or SD event were excluded

## Discussion

Epileptic animal models (Mellanby, George et al. 1977, Cavalheiro, Riche and Le Gal La Salle 1982, Lothman, Bertram et al. 1990, Cavalheiro, Leite et al. 1991, Stafstrom, Thompson and Holmes 1992, Bertram and Cornett 1994, JEFFERYS 2006) characterized by recurrent spontaneous seizures closely mimic human epilepsy and are therefore more valuable for studying seizure mechanisms, associated comorbidities, and treatment development (Rusina, Bernard and Williamson 2021). However, capturing these spontaneous seizures requires long-term continuous EEG/video recordings which are typically resource-intensive (Bertram, Williamson et al. 1997). Our EMU can provide chronic, 24/7 continuous biopotential and video recordings on multiple animals in a low-cost, high-fidelity, and time-synchronized way. Each rig within the EMU fits commercial racks and takes advantage of auto-clave-ready cages without extra components and can be duplicated for under $1000. We demonstrated LFP, ECoG, and ECG recordings on multiple freely moving rats without interruption even during animal handling. Data from different rats was also time synchronized and stored in one NAS for offline analysis. The high-fidelity, DC-sensitive recordings permitted us to capture spontaneous seizures and seizure-associated SDs.

In the TeTX rat model, SD is associated with more than a third of all seizures. Furthermore, as shown in Figure 10, following ictal onset, seizures that evolve into SD undergo a silent period, followed by recovery after-discharges, i.e., spreading convulsions. DC-sensitive recordings offer deeper insights into this seizure event by revealing the SD occurrence and confirming that it is indeed not a separate event but rather a part of the same seizure. There are many peri and post-ictal phenomena whose underlying mechanisms may be linked to seizure-associated SD, including post-ictal generalized EEG suppression (PGES), postictal amnesia, loss of consciousness, memory deficits, headaches, and death. PGES has been extensively investigated and linked to post-ictal cardio-respiratory dysregulation and impaired cognition (Bateman, Li and Seyal 2008, Lhatoo, Faulkner et al. 2010, Seyal, Pascual et al. 2011, Seyal, Hardin and Bateman 2012). There are hypotheses that SD might be the underlying phenomenon of PGES (Rajakulendran and Nashef 2015).

Sudden unexplained death in epilepsy (SUDEP) affects approximately 1 in 1000 adults with epilepsy per year (Thurman, Hesdorffer and French 2014). Recent studies (Aiba and Noebels 2015) implicate brainstem SD as a mechanism of SUDEP and have been supported by recent *in-vivo* recording and imaging studies (Loonen, Jansen et al. 2019). Clinically, seizure-associated SD has been proposed as a biomarker of imminent SUDEP (Lhatoo, Noebels et al. 2015). Our EMU is suitable for SUDEP animal model studies.

The EMU subsystems can be used individually or in combination with several for different applications. Acceleration parameters have been utilized for the last six decades to investigate pathology in both human and animal models of traumatic brain injury (TBI), design safety equipment, and develop injury thresholds (Mayer, Ling et al. 2021).

Acceleration measures have also been used to identify behavioral modes (2012, Chen, Gu et al. 2019, McNamara, Grillakis et al. 2020, Mayer, Ling et al. 2021, Maekawa, Sakitani et al. 2022). The recording system provides a 3-axis head-acceleration measurement in our application which enabled us to monitor the head-bobbing and freezing behavior and full-body convulsions during seizure and post-ictal periods. The commutator and isolation system can be used for experiments on freely moving animals. In many rodent animal epilepsy studies, animals are frequently in an anesthetized state. However, anesthesia may alter the normal brain function directly or indirectly. Compared with head-fixed or anesthetized mice, recording neural activity in awake freely moving mice is a powerful and flexible technique for dissecting the neural circuit mechanisms underlying pathological behavior (Harris, Golder and Likhtik 2017). However, recordings from freely moving animals involve a cable connection for data transmission, restricting the animal’s behavior. Wireless recording systems have been applied in freely moving animal studies (Kramer and Kinter 2003, Kadam, White et al. 2010, Zayachkivsky, Lehmkuhle and Dudek 2015, Lundt, Wormuth et al. 2016). However, these telemetry systems face two issues for long-term (weeks to months) continuous monitoring work. First, the battery life is usually hours, and cannot finish chronic continuous recordings to capture spontaneous seizures. Second, the data transmission rate is low and the connection is unstable. In our commutator and isolation system, the commutator supports a standalone operation without additional supporting devices like counterbalanced bars or ball bearings. Unlike commercial products requiring specific cable connections, it provides a direct USB connection. The isolation makes the whole system human-compatible and is suitable for brain computer interface (BCI) applications. We demonstrated, in our TeTX model, that development of abnormal brain-to-heart coupling prior to the first convulsive seizure may be adapted to forecast seizure clusters (seizure clusters example shown in Suppl. Figure 1) with long seizure-free intervals between them. The cortically-induced disturbances of cardiac rhythm maybe responsible for making the network more susceptible to more seizures and vulnerable to their effects (Lathers, Schraeder and Weiner 1987, Schuele, Bermeo et al. 2007). Electrochemical methods comprise a collection of extremely useful measurement tools for neuroscience (Borland and Michael 2007). Electrochemical measurements in brain tissue, especially different voltammetric measures provide important complementary signals, e.g., tissue oxygenation, along with behavioral, electrophysiological, and optical measurements. Tissue oxygenation measured using an electrochemical way is the ‘gold standard’ (Stone, Brown et al. 1993, Vaupel, Hockel and Mayer 2007). *In-vivo* electrochemistry has grown steadily over the past several decades. Adams’ early slow-scan voltammetry experiments, which made use of carbon-paste microelectrodes, have given way to high temporal resolution voltammetry measurements at micron- and sub-micron-scale electrodes (Johnson 2013). Different voltammetric and amperometric techniques have been combined with electrophysiology and enzyme-mediated biosensors (Wilson and Johnson 2008), thereby expanding the tools available to investigators (Johnson 2013). *In-vivo* electrochemistry has also been applied for neurotransmitter sensing. Using FSCV, norepinephrine and dopamine sensing (Park, Takmakov and Wightman 2011) has been done in rats and norepinephrine sensing has been reviewed in detail by (Park, Bhimani and Bass 2018). Long-term *in-vivo* electrochemistry enables real-time monitoring and measurement of brain metabolites (Lowry, Griffin et al. 2010).

Generally, an electrochemical instrumentation (EI) consists of a potentiostat (or galvanostat), for applying a controlled potential (or current) on an electrode along with a function generator, to produce the desired perturbation, and a recording system for measuring the response current (or potential). Using the built-in EI in the EMU, we demonstrated an *in-vitro* oxygen-sensing electrode calibration process and robust *in-vivo* oxygen sensing using CPA on freely moving animals coupled with electrophysiology recordings. Although CPA is known to yield local tissue oxygenation and has been widely used for in vivo oxygen sensing. Studies using the CPA for oxygen sensing either assume that the oxygen diffusion coefficient is constant or are incapable of decoupling the effects of local oxygen concentration and the oxygen diffusion coefficient on the CPA measure (Subczynski and Swartz 2005). A further innovative way to do tissue oxygenation measure is needed for scenarios, e.g. during SD event, when the oxygen diffusion coefficient changes.

Considering the built-in function generator, new oxygenation measure is feasible during peri-seizure and peri-SD events. Moreover, our EMU shows potential for studying neurovascular coupling and metabolic dynamics in freely moving animals by combining *in-vivo* pO_2_ measurements with electrophysiology and neural imagings. It could be particularly valuable for investigating pathological states such as seizures and SD that need long-term, high-fidelity, continuous recordings.

## Author Contributions

JL and BJG designed and constructed the hardware, designed the research, analyzed the data, and wrote the paper.

## Acknowledgement

We thank the participation of Fatemeh Bahari, John Kimbugwe, and Carlos Curay who served in implanting the animals and collecting data.

---

**Supplementary Figure 1. Example of seizure clusters.** Generated from Loki011. The upper panel shows band-filtered ECoG and hippocampal/ thalamus LFP measurements, and the lower panel displays low-pass filtered differential hippocampal/ thalamus LFP measurements. The ictal period is marked in yellow. The lower pane also shows DC-band recordings of seizures and associated spreading depolarizations (SD), with triangles indicating the SD propagation direction.

