## Supplementary figures and images for "A DC-sensitive video/electrophysiology monitoring unit for long-term continuous study of seizures and seizure-associated spreading depolarization in a rat model"

### Supplemental Figure 1

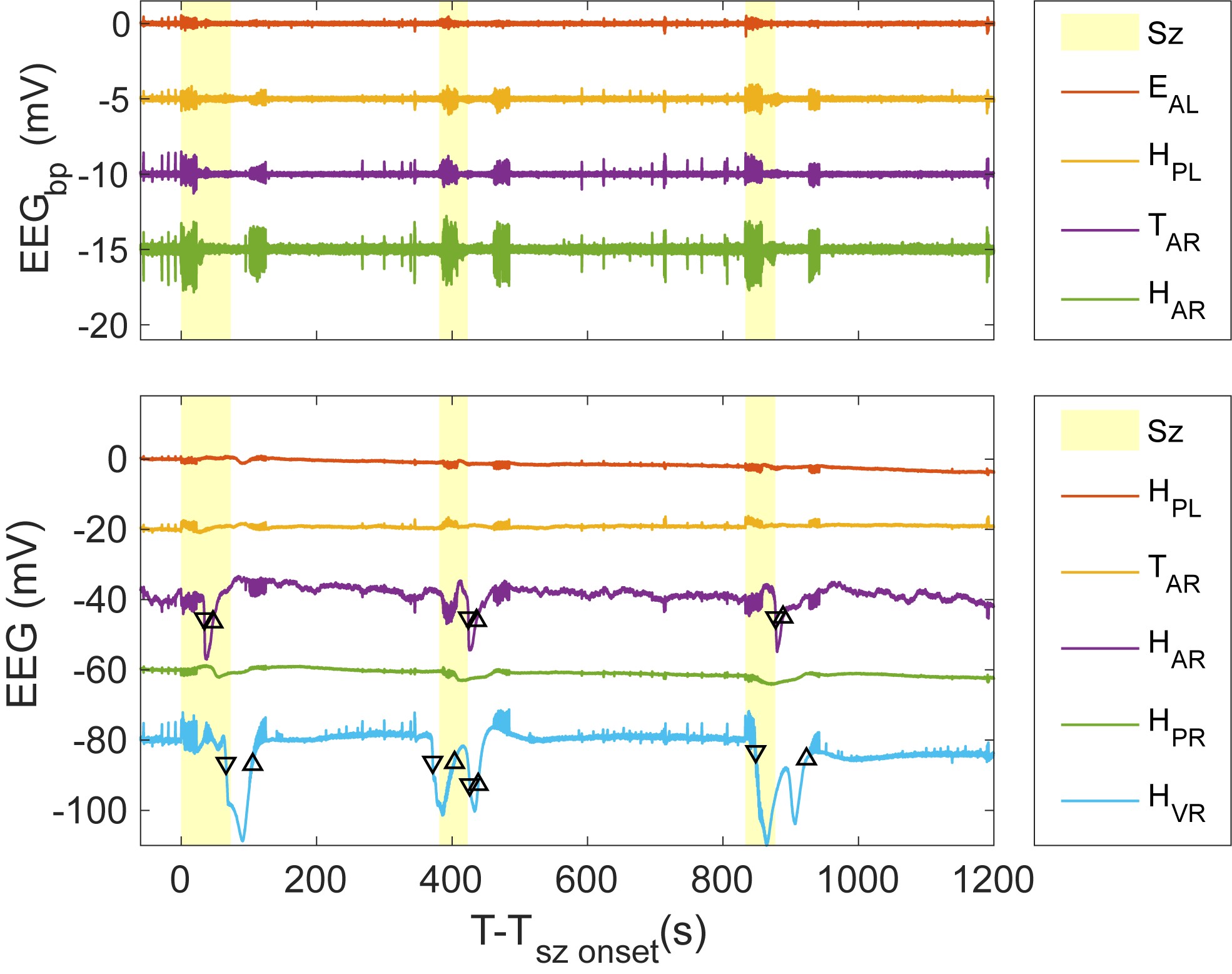
